# Characterization of the untranslated region of lymphocytic choriomeningitis virus S segment

**DOI:** 10.1101/653808

**Authors:** Satoshi Taniguchi, Tomoki Yoshikawa, Masayuki Shimojima, Shuetsu Fukushi, Takeshi Kurosu, Hideki Tani, Aiko Fukuma, Fumihiro Kato, Eri Nakayama, Takahiro Maeki, Shigeru Tajima, Chang-Kweng Lim, Hideki Ebihara, Shigeru Kyuwa, Shigeru Morikawa, Masayuki Saijo

## Abstract

Lymphocytic choriomeningitis virus (LCMV) is a prototypic arenavirus. The viral genome consists of two RNA segments, L and S. The 5’- and 3’-termini of both L and S segments are highly conserved among arenaviruses. These regions consist of 19 complementary base pairs and are essential for viral genome replication and transcription. In addition to these 19 nucleotides in the 5’- and 3’-termini, there are untranslated regions (UTRs) composed of 58 and 41 nucleotide residues in the 5’ and 3’ UTRs, respectively, in the LCMV S segment. Their functional roles, however, have yet to be elucidated. In this study, a reverse genetics and a minigenome system for the LCMV strain WE were established and used to analyze the function of these regions. The results obtained from these analyses, plus RNA secondary structure prediction, revealed that not only these 19 nucleotides but also the 20th–40th and 20th–38th nucleotides located downstream of the 19 nucleotides in the 5’- and 3’-termini, respectively, are heavily involved in viral genome replication and transcription. Furthermore, the introduction of mutations in these regions depressed viral propagation *in vitro* and enhanced attenuation *in vivo*. Conversely, recombinant LCMVs (rLCMVs), which had various deletions in the other UTRs, propagated as well as wild-type LCMV *in vitro* but were attenuated *in vivo*. Most mice previously infected with rLCMVs with mutated UTRs, when further infected with a lethal dose of wild-type LCMV, survived. These results suggest that rLCMVs with mutated UTRs could be candidates for an LCMV vaccine.

**IMPORTANCE:** The function of untranslated regions (UTRs) of the arenavirus genome has not well been studied except for the 19 nucleotides of the 5’- and 3’-termini. In this study the function of the UTRs of the LCMV S segment was analyzed. It was found that not only the 19 nucleotides of the 5’- and 3’-termini but also the 20th–40th and 20th–38th nucleotides located downstream of the 19 nucleotides in the 5’- and 3’-termini, respectively, were involved in viral genome replication and transcription. Furthermore, other UTRs in the S segment were involved in virulence *in vivo*. The introduction of mutations to these regions makes it possible to establish attenuated LCMV and potentially develop LCMV vaccine candidates.

## INTRODUCTION

Lymphocytic choriomeningitis virus (LCMV) belongs to the genus *Mammarenavirus*, family *Arenaviridae*. There are two groups of *Mammarenavirus*, New World and Old World arenaviruses (1). LCMV belongs to the Old World arenavirus group, as does Lassa virus (LASV), the causative agent of Lassa fever. LCMV can infect humans, causing flu-like fever, nausea, neck stiffness, headache, and occasionally photophobia. Severe cases can lead to meningitis and encephalitis (2–6). The natural reservoir of LCMV is reported to be the house mouse (*Mus musculus*). Humans can be infected with LCMV if they are exposed to the body fluids of infected mice.

Since LCMV was first isolated in 1933, it has been one of the most widely used model systems with which to study immunology, persistent infection, and pathogenesis relating to viruses (7–9). Furthermore, LCMV is the prototype arenavirus, and the LCMV strain Armstrong (LCMV-ARM), a laboratory strain that causes acute neurotropic infection in mice, has been commonly used in these studies. Minigenome and reverse genetics systems based on LCMV-ARM have been developed and are powerful tools used for the study of LCMV-ARM pathogenesis and propagation mechanisms (10–17). However, molecular biological analyses based on other strains of LCMV have not previously been performed.

Infection with different strains of LCMV causes different manifestations of disease in various animal models (18–22). For example, the LCMV-ARM and LCMV strain WE (LCMV-WE) cause neurotropic and viscerotropic symptoms in mice, respectively. LCMV-ARM infection causes no symptoms in rhesus macaques, while LCMV-WE causes hemorrhagic fever, hepatic damage, and meningitis in this species (21, 23–27). The infection of non-human primates with LCMV-WE is considered a suitable animal model for giving insights into Lassa fever in humans. Despite its uniqueness and importance there have been few basic tools developed, such as a reverse genetics system, to analyze the virologic characteristics of LCMV-WE.

The genome of LCMV consists of two negative-sense single-stranded RNA segments, designated S and L. The S segment, which is approximately 3.4 kilobases (kb) in length, encodes a viral glycoprotein precursor (GPC) and a nucleoprotein (NP), while the L segment, which is approximately 7.2 kb long, encodes a viral RNA-dependent RNA polymerase (L) and a polypeptide that contains a small zinc finger-domain (Z) (1). Each segment has an ambisense coding strategy, encoding two proteins in opposite orientations, separated by an intergenic region (IGR) that folds into a predictable and stable secondary structure. Recently, it was reported that replacement of the L segment IGR with the S segment IGR resulted in highly attenuated LCMV *in vivo* and induced protective immunity against a lethal challenge with wild-type LCMV (wtLCMV) (28).

Untranslated regions (UTRs) are located at the termini of each genome segment, and the 19 nucleotides of the 5’- and 3’-termini of both S and L segments are reported as being essential regions for viral transcription and replication (29, 30). These regions in the S and L segments are predicted to comprise complementary base pairs and are recognized as promoters of L-polymerase-driven viral genome transcription and replication (29, 30). However, the functional roles in virulence and propagation of other regions within the UTRs have yet to be elucidated. To investigate the roles of these regions, we developed reverse genetics and minigenome systems for LCMV-WE and predicted the secondary structures of the RNA sequences of the 5’- and 3’-terminal UTRs of the S segment. Based on the results of our RNA secondary structure prediction, several variants of infectious clones lacking UTRs, except for the 19 nucleotides of the 5’- and 3’-termini of the S segment, were generated and the viral propagation efficiency of the infectious clones was evaluated. The virulence of these infectious clone variants in mice was also assessed. Furthermore, we generated variants of minigenome RNAs lacking the UTRs of the S segment and elucidated their efficiency in viral genome replication, transcription, and packaging of virus-like particles (VLPs).

## RESULTS

### Design of plasmids for recombinant LCMV with mutations in the S segment UTRs and prediction of their RNA secondary structure

The results of our RNA secondary structure predictions, made using the CENTROIDFOLD server, are shown in Fig. 1 and 2. RNA produced from pRF-WE-SRG (i.e., LCMV genome segment S RNA) formed panhandle structures of 45 and 42 nucleotide (nt) at the 5’- and 3’-termini, respectively (Fig. 1A). pRF-WE-SRG was a plasmid prepared for the LCMV-WE reverse genetics system, which coded the entire LCMV-WE S segment. In the terminal panhandle structures, the 28th–33rd and the 26th–31st nt at the 5’- and 3’-termini, respectively, formed base pairs with the highest base-pairing probability in addition to the 19 base pairs at the termini. Focusing on the panhandle structure, in particular the base pairs comprising the 28th– 33rd nt at the 5’-terminus and the 26th–31st nt at the 3’-terminus, we designed pRF-WE-SRGs with various mutations, which could generate various mutated LCMV S segment RNAs (Table 1 and 2). The results of the prediction of RNA secondary structures of RNA products derived from pRF-WE-SRG-5UTRΔ20-40, pRF-WE-SRG-5UTRΔ41-60, pRF-WE-SRG-5UTRΔ60-77, pRF-WE-SRG-3UTRΔ20-38, and pRF-WE-SRG-3UTRΔ39-60 are shown in Fig. 1B to 1F. The RNAs produced from pRF-WE-SRG-5UTRΔ41-60, pRF-WE-SRG-5UTRΔ60-77, and pRF-WE-SRG-3UTRΔ39-60 formed panhandle structures of more than 36 and 33 nt at the 5’-and 3’-termini, respectively. The base pairs comprising the 28th–33rd nt at the 5’-terminus and the 26th– 31st nt at the 3’-terminus showed the highest base-pairing probability. The RNAs produced from pRF-WE-SRG-5UTRΔ20-40 and pRF-WE-SRG-3UTRΔ20-38 did not form terminal panhandle structures with the exception of the 19 base pairs in the termini.

**FIG 1.**
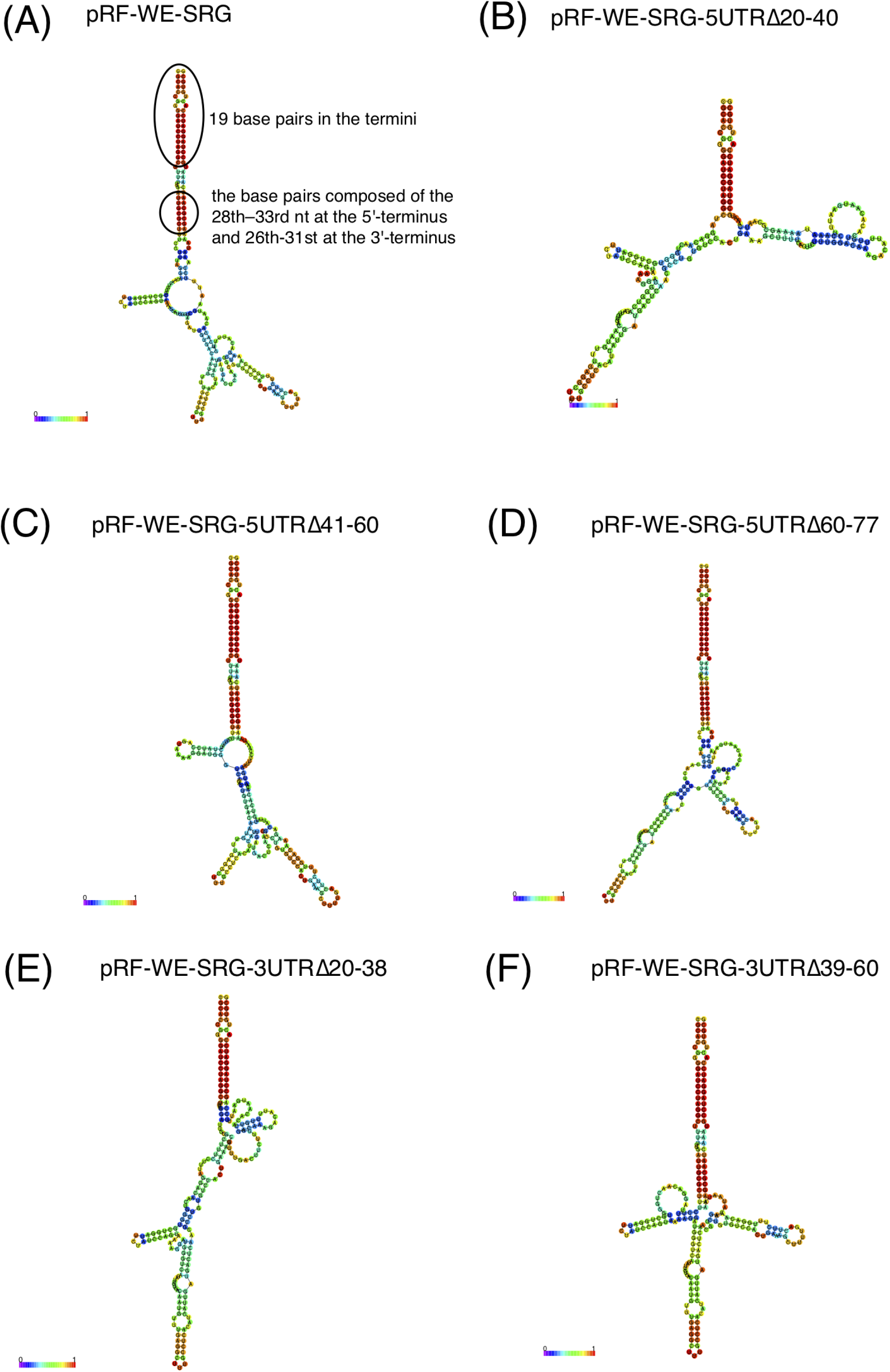
Prediction of RNA secondary structures of LCMV strain WE (LCMV-WE) S segment UTR and its various mutated UTRs which have 18–22 nt deletions in the 5’- or 3’-termini. (A) Predicted RNA secondary structure of LCMV-WE S segment UTR derived from pRF-WE-SRG. The location of the 19 base pairs in the termini and the base pairs comprising the 28th–33rd nt in the 5’-terminus and the 26th–31st nt in the 3’-terminus are shown (B to F). Predicted RNA secondary structures of LCMV-WE S segment UTRs which have 18–22 nt deletions in the 5’- or 3’-terminal UTRs. These RNAs were derived from pRF-WE-SRG-5UTRΔ20-40 (B), pRF-WE-SRG-5UTRΔ41-60 (C), pRF-WE-SRG-5UTRΔ60-77 (D), pRF-WE-SRG-3UTRΔ20-38 (E), and pRF-WE-SRG-3UTRΔ39-60 (F). RNA sequences of LCMV-WE S segment genome 5’-terminal and 3’-terminal UTRs (or various mutated UTRs) and 50 nt of ORF regional RNA sequences that were directly downstream of the UTRs were linked, sent to the CENTROIDFOLD server, and analyzed using the CONTRAfold model (weight of base pairs: 2^2). Each predicted base pair is colored with heat-color gradation from blue to red, corresponding to the base-pairing probability from 0 to 1. Detailed information about pRF-WE-SRG and the various mutated pRF-WE-SRGs is given in Table 1 and 2.

**FIG 2.**
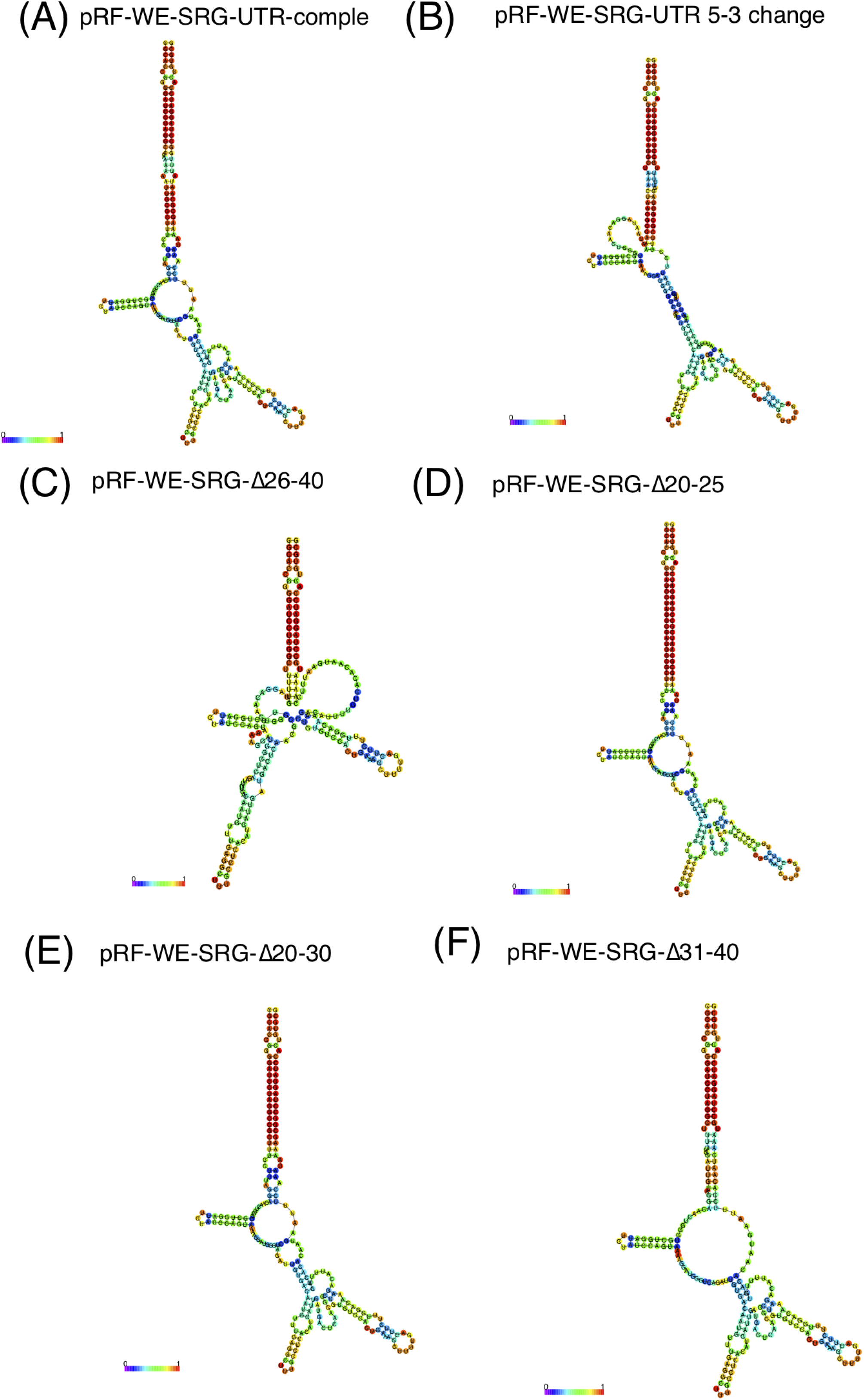
Prediction of RNA secondary structures of mutated LCMV strain WE (LCMV-WE) S segment UTRs which have deletions or mutations in the 20th–40th nt in the 5’-terminus or 20th– 38th nt in the 3’-terminus (A to F). These RNAs were derived from pRF-WE-SRG-UTR-comple (A), pRF-WE-SRG-UTR 5-3 change (B), pRF-WE-SRG-Δ20-30 (C), pRF-WE-SRG-Δ26-40 (D), pRF-WE-SRG-Δ31-40 (E), and pRF-WE-SRG-Δ20-25 (F). RNA sequences of the mutated LCMV-WE S segment genome 5’-terminal and 3’-terminal UTRs and 50 nt of ORF regional RNA sequences that were directly downstream of the UTRs were linked, sent to the CENTROIDFOLD server, and analyzed using the CONTRAfold model (weight of base pairs: 2^2). Each predicted base pair is colored with heat-color gradation from blue to red, corresponding to the base-pairing probability from 0 to 1. Detailed information about the various mutated pRF-WE-SRGs is given in Table 1 and 2.

**Table 1.**
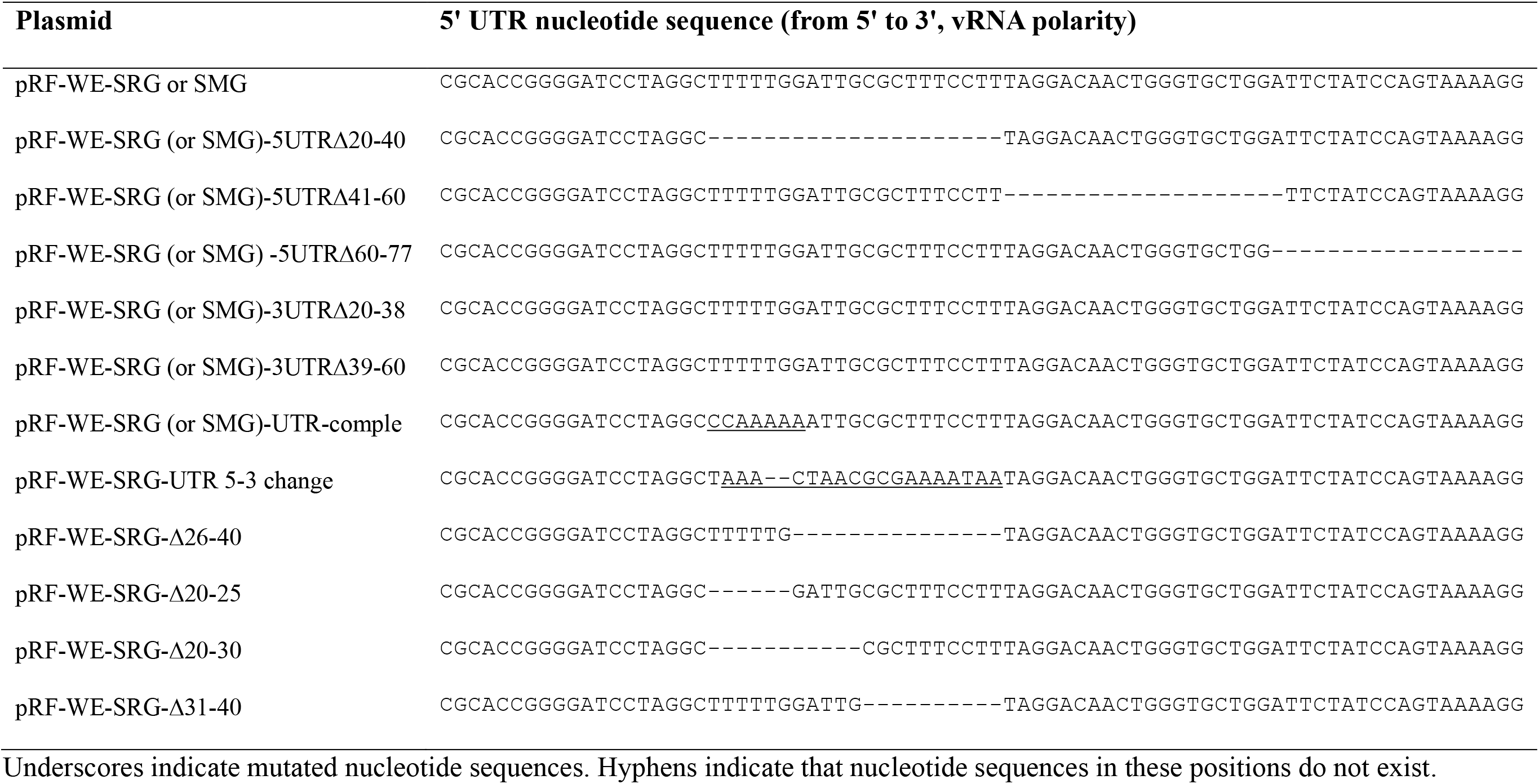
The 5’ nucleotide sequences of untranslated regions (UTRs) of the plasmids for the reverse genetics and minigenome systems.

**Table 2.**
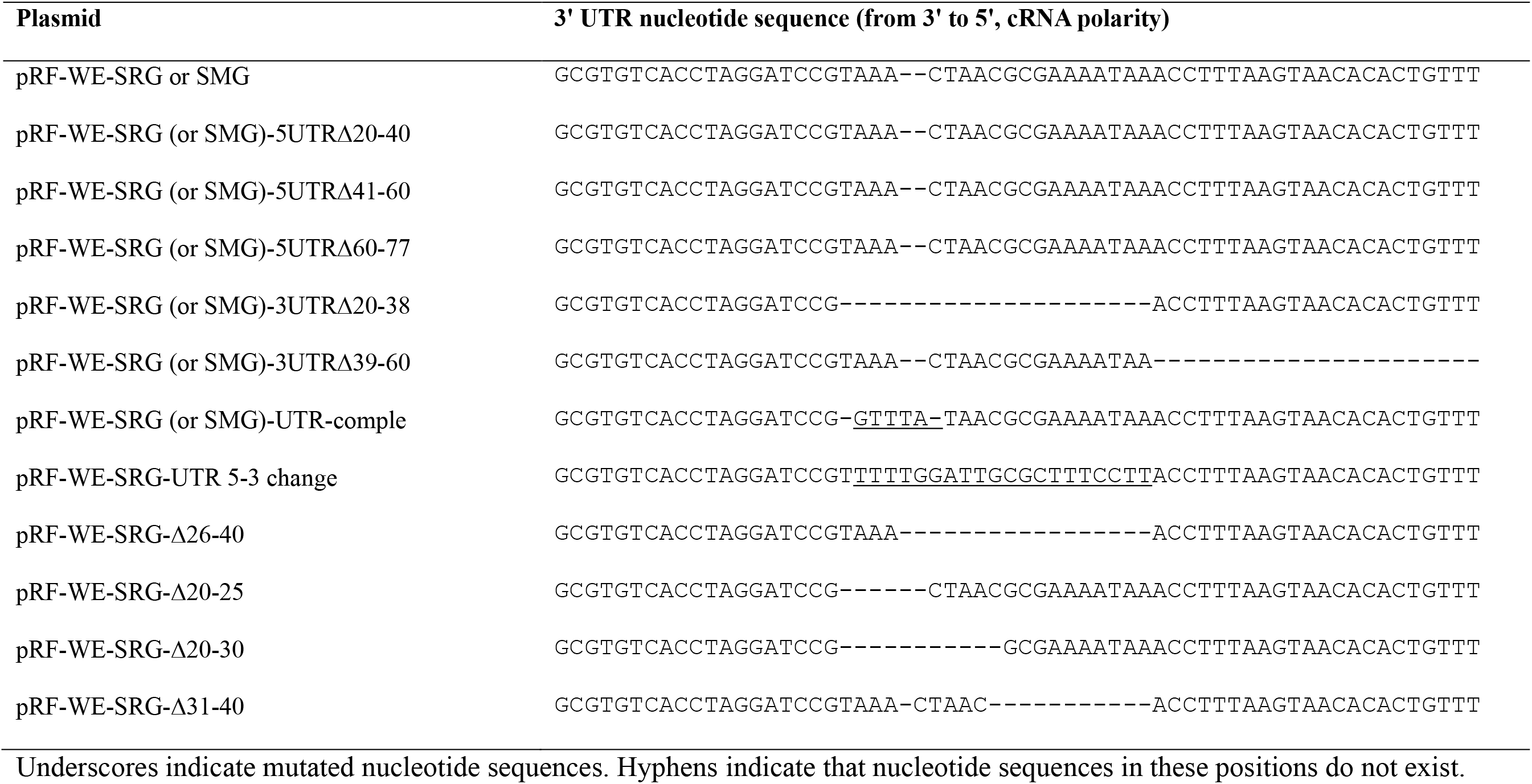
The 3’ nucleotide sequences of untranslated regions (UTRs) of the plasmids for the reverse genetics and minigenome systems.

To further investigate the effect of the base pairs comprising the 28th–33rd nt at the 5’-terminus and the 26th–31st nt at the 3’-terminus and their peripheral regions, various plasmids with mutations or deletions in the regions of interest (20th–40th nt at the 5’-terminus and 20th– 38th nt at the 3’-terminus) were generated. The results of RNA secondary structure prediction of the RNA products of pRF-WE-SRG-UTR-comple, pRF-WE-SRG-UTR 5-3 change, pRF-WE-SRG-Δ26-40, pRF-WE-SRG-Δ20-25, pRF-WE-SRG-Δ20-30, and pRF-WE-SRG-Δ31-40 are shown in Fig. 2. A summary of the results of the RNA secondary structure predictions is shown in Table 3. Briefly, the RNAs produced from pRF-WE-SRG-UTR-comple, pRF-WE-SRG-UTR 5-3 change, pRF-WE-SRG-Δ20-25 and pRF-WE-SRG-Δ20-30 formed panhandle structures with 5’- and 3’-termini which were composed of the base pairs with high base-pairing probability, in addition to the 19 base pairs in the termini. On the other hand, the RNAs produced from pRF-WE-SRG-Δ26-40 and pRF-WE-SRG-Δ31-40 formed panhandle structures with 5’- and 3’-termini but there were no base pairs with high base-pairing probability except for the 19 base pairs in the termini.

**Table 3.** Summary of the results of RNA secondary structure prediction of the LCMV S segment RNA produced from each plasmid.

| Plasmid | Number of nucleotides forming a panhandle structure at the termini |  | Base pairs with high base-pairing probability, except for the 19 base pairs at the termini |  |
| --- | --- | --- | --- | --- |
|  | At the 5'-terminus | At the 3'-terminus | At the 5'-terminus | At the 3'-terminus |
| pRF-WE-SRG | 45 nt | 42 nt | 28th–33rd nt | 26th–31st nt |
| pRF-WE-SRG-5UTRΔ20-40 | 18 nt | 18 nt | none | none |
| pRF-WE-SRG-5UTRΔ41-60 | 36 nt | 33 nt | 28th–33rd nt | 26th–31st nt |
| pRF-WE-SRG-5UTRΔ60-77 | 45 nt | 42 nt | 28th–33rd nt | 26th–31st nt |
| pRF-WE-SRG-3UTRΔ20-38 | 27 nt | 23 nt | none | none |
| pRF-WE-SRG-3UTRΔ39-60 | 36 nt | 33 nt | 28th–33rd nt | 26th–31st nt |
| pRF-WE-SRG-UTR-comple | 45 nt | 42 nt | 28th–33rd nt | 26th–31st nt |
| pRF-WE-SRG-UTR 5-3 change | 34 nt | 36 nt | 26th–31st nt | 28th–33rd nt |
| pRF-WE-SRG-Δ26-40 | 25 nt | 25 nt | none | none |
| pRF-WE-SRG-Δ20-25 | 39 nt | 39 nt | 20th–27th nt | 20th–27th nt |
| pRF-WE-SRG-Δ20-30 | 34 nt | 33 nt | 20th–22nd nt | 20th–22nd nt |
| pRF-WE-SRG-Δ31-40 | 35 nt | 32 nt | none | none |

### Rescue and characterization of recombinant LCMVs

First, recombinant non-mutated wild-type LCMV (rwtLCMV) was successfully generated using pRF-WE-SRG. There was no significant difference in viral growth kinetics in Vero cells between wild-type LCMV (wtLCMV) and rwtLCMV (*P* = 0.0723) (Fig. 3A). Next, we tried to generate recombinant LCMVs (rLCMVs) from pRF-WE-SRG-5UTRΔ20-40, pRF-WE-SRG-5UTRΔ41-60, pRF-WE-SRG-5UTRΔ60-77, pRF-WE-SRG-3UTRΔ20-38, and pRF-WE-SRG-3UTRΔ39-60. The generation of rLCMVs using pRF-WE-SRG-5UTRΔ41-60 (rLCMV-5UTRΔ41-60), pRF-WE-SRG-5UTRΔ60-77 (rLCMV-5UTRΔ60-77), pRF-WE-SRG-3UTRΔ39-60 (rLCMV-3UTRΔ39-60) was confirmed by immunofluorescent assay (IFA). The generation of rLCMVs using pRF-WE-SRG-5UTRΔ20-40 and pRF-WE-SRG-3UTRΔ20-38 could not be confirmed. The viral growth kinetics of rLCMV-5UTRΔ41-60, rLCMV-5UTRΔ60-77, and rLCMV-3UTRΔ39-60 in Vero cells were compared with rwtLCMV or wtLCMV, the result being that all LCMVs propagated efficiently in Vero cells with titers up to 1.0 × 10^6^–10^7^ focus-forming units (FFU)/ml at 72 hours post-infection (Fig. 3B). No significant differences in viral growth efficiency in Vero cells were observed among rwtLCMV, rLCMV-5UTRΔ41-60, or rLCMV-5UTRΔ60-77 (*P* = 0.4698 and 0.3250; total variation, which consists of the sum of the squares of the differences of each mean with the grand mean: 0.06001% and 0.6165%, respectively). A significant difference in viral growth efficiency in Vero cells was observed between rwtLCMV and rLCMV-3UTRΔ39-60 (*P* = 0.0001; total variation = 5.941%), but no significant difference in viral growth efficiency in Vero cells was observed between wtLCMV and rLCMV-3UTRΔ39-60 (*P* = 0.6755; total variation = 0.04217%).

**FIG 3.**
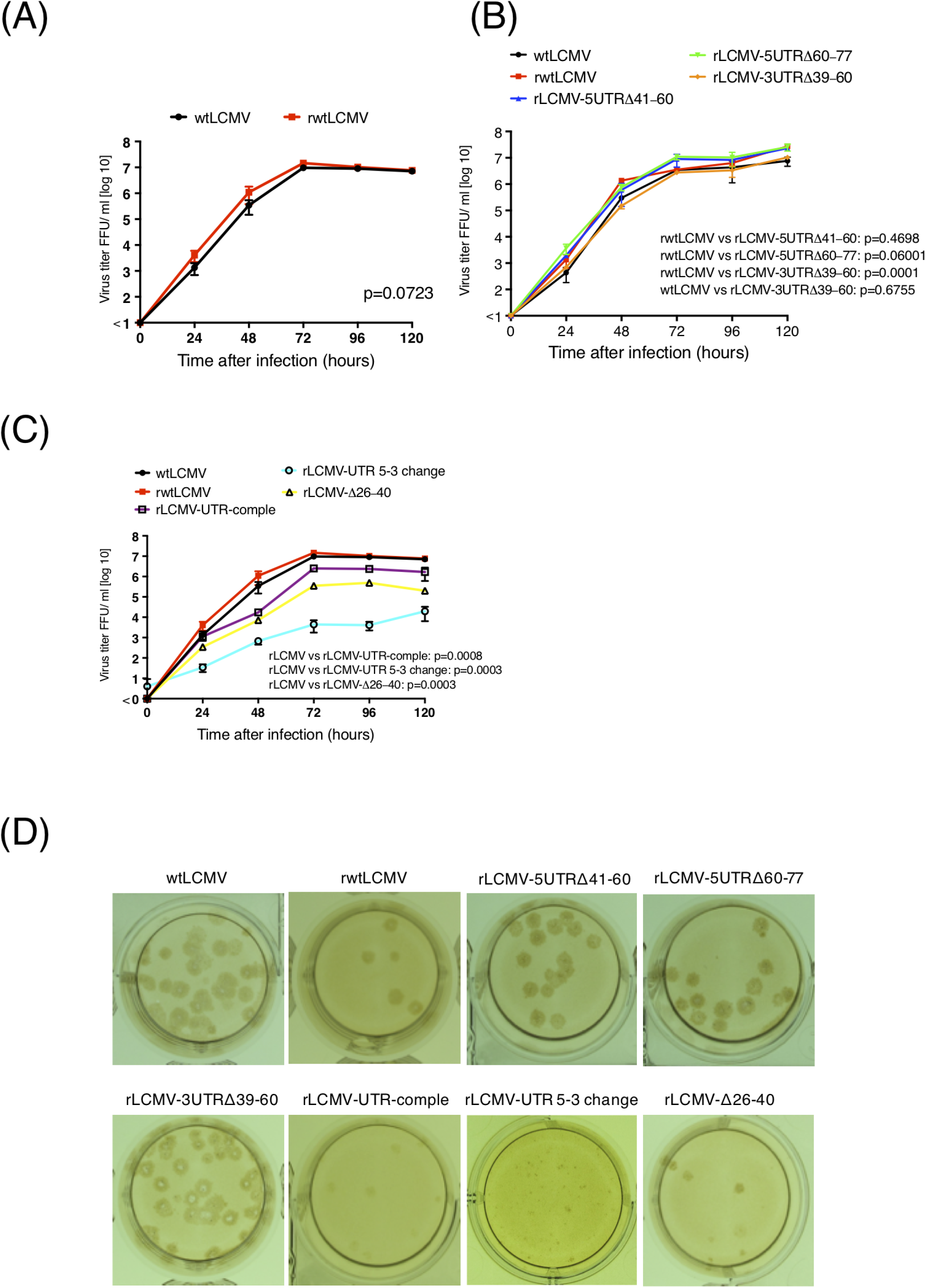
Viral growth properties in Vero cells. Confluent monolayers of Vero cells were infected with wtLCMV or rLCMVs at an moi of 0.01 per cell. Cells were washed three times with DMEM-2%FBS after a one-hour adsorption period, and 1 ml DMEM-2%FBS was added to each well. Supernatant samples were collected at 0, 24, 48, 72, 96, and 120 hours post-infection. The supernatants were centrifuged at 8,000 rpm for 5 min to remove cell debris and stored at −80°C. The infectious dose was measured using a viral immunofocus assay. Viral growth curves of LCMVs were statistically analyzed using two-way ANOVA. (A) Viral growth properties of wild-type LCMV (wtLCMV) and recombinant non-mutated wild-type LCMV (rwtLCMV) in Vero cells were compared. (B) Viral growth kinetics of rLCMV-5UTRΔ41-60, rLCMV-5UTRΔ60-77, and rLCMV-3UTRΔ39-60 in Vero cells were compared with wtLCMV and rwtLCMV. (C) Viral growth kinetics of rLCMV-UTR-comple, rLCMV-UTR 5-3 change, and rLCMV-Δ26-40 in Vero cells were compared with wtLCMV and rwtLCMV. (D) Focus morphology of wtLCMV and rLCMVs in Vero cells. The focus morphologies of wtLCMV and rLCMVs generated in this study are shown. Error bars in FIG 3A, B, and C indicate standard deviations.

rLCMVs with mutations in the UTRs of interest (the 20th–40th nt at the 5’-terminus and the 20th-38th nt at the 3’-terminus) were generated from pRF-WE-SRG-UTR-comple, pRF-WE-SRG-UTR 5-3 change, pRF-WE-SRG-Δ26-40, pRF-WE-SRG-Δ20-25, pRF-WE-SRG-Δ20-30, and pRF-WE-SRG-Δ31-40 (Table 4). The generation of rLCMVs using pRF-WE-SRG-UTR-comple (rLCMV-UTR-comple), pRF-WE-SRG-UTR 5-3 change (rLCMV-UTR 5-3 change), and pRF-WE-SRG-Δ26-40 (rLCMV-Δ26-40) was confirmed. Because the propagation efficiency of rLCMV-UTR 5-3 change and rLCMV-Δ26-40 was low, they were further passaged in Vero cells before use. The generation of rLCMVs using pRF-WE-SRG-Δ20-25 (rLCMV-Δ20-25), pRF-WE-SRG-Δ20-30 (rLCMV-Δ20-30), and pRF-WE-SRG-Δ31-40 (rLCMV-Δ31-40) was not confirmed. The viral growth capacity of rLCMV-UTR-comple, rLCMV-UTR 5-3 change, and rLCMV-Δ26-40 in Vero cells was significantly less than that of rwtLCMV (Fig. 3C, *P* = 0.0008, 0.0003, and 0.0003; total variation = 22.55%, 31.86%, and 30.51%, respectively).

**Table 4.**
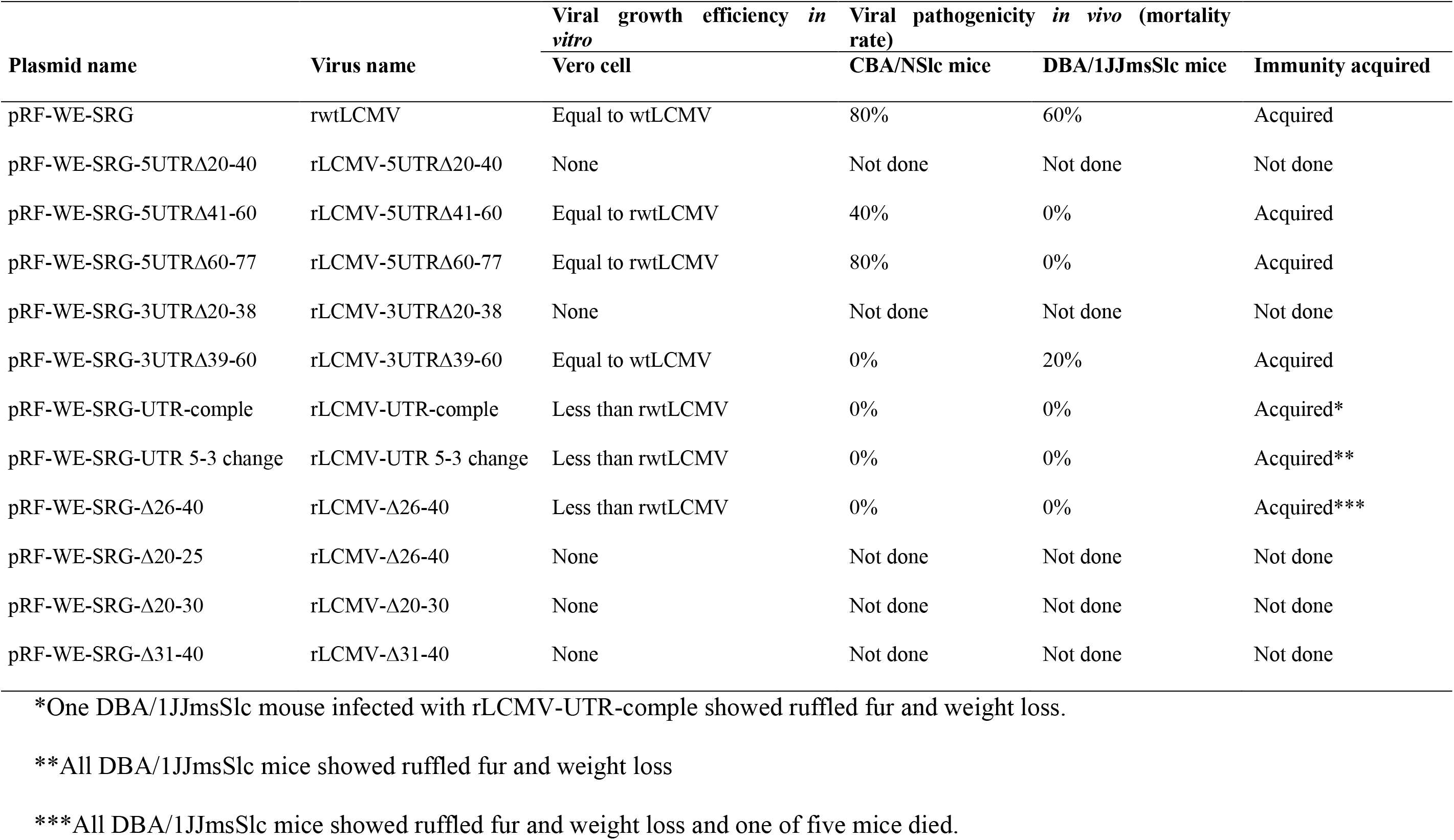
Summary of viral growth efficiency *in vitro* and pathogenicity *in vivo* of each recombinant LCMV generated from each plasmid.

The focus morphologies of wtLCMV and rLCMVs generated in this study are shown in Fig. 3D. The size of foci was concordant with viral growth-capacity features. The focus sizes of rLCMV-5UTRΔ41-60, rLCMV-5UTRΔ60-77, and rLCMV-3UTRΔ39-60 were equivalent to those of wtLCMV and rwtLCMV. The focus sizes of rLCMV-UTR-comple and rLCMV-Δ26–40 were smaller than those of wtLCMV and rwtLCMV. The focus size of rLCMV-UTR 5-3 change was the smallest among them.

### Pathogenicity of rLCMVs in mice

To investigate the virulence of the rLCMVs (rwtLCMV, rLCMV-5UTRΔ41-60, rLCMV-5UTRΔ60-77, rLCMV-3UTRΔ39-60, rLCMV-UTR-comple, rLCMV-UTR 5-3 change, and rLCMV-Δ26-40), CBA/NSlc and DBA/1JJmsSlc mice were infected intraperitoneally (i.p.) with wtLCMV or the generated rLCMVs at 1.0 × 10^2^ FFU/head.

CBA/NSlc mice infected with wtLCMV (wtLCMV-CBA/NSlc mice), rwtLCMV (rwtLCMV-CBA/NSlc mice), rLCMV-5UTRΔ41-60 (rLCMV-5UTRΔ41-60-CBA/NSlc mice), or rLCMV-5UTRΔ60-77 (rLCMV-5UTRΔ60-77-CBA/NSlc mice) showed several clinical signs of infection, such as ruffled fur and limb tremors at 7 days post-infection (d.p.i.), and weight loss at 8 d.p.i. (Fig. 4A). All wtLCMV-CBA/NSlc mice, 4 out of 5 rwtLCMV-CBA/NSlc mice, 4 out of 5 rLCMV-5UTRΔ60-77-CBA/NSlc mice, and 2 out of 5 rLCMV-5UTRΔ41-60-CBA/NSlc mice died within 20 d.p.i. (Fig. 4B and Table 4). Significant differences in survival rates were not observed between rwtLCMV and rLCMV-5UTRΔ60-77 (*P* = 0.3997) or between rwtLCMV and rLCMV-5UTRΔ41-60 (*P* = 0.3148). Hazard ratios for rwtLCMV / rLCMV-5UTRΔ60-77 and rwtLCMV / rLCMV-5UTRΔ41-60 were 1.517 and 2.266, respectively. On the other hand, all CBA/NSlc mice infected with either rLCMV-3UTRΔ39-60 (rLCMV-3UTRΔ39-60-CBA/NSlc mice), rLCMV-UTR-comple (rLCMV-UTR-comple-CBA/NSlc mice), rLCMV-UTR 5-3 change (rLCMV-UTR 5-3 change -CBA/NSlc mice), or rLCMV-Δ26-40 (rLCMV-Δ26-40-CBA/NSlc mice) showed no clinical signs and survived. Significant differences in survival rates were observed between rwtLCMV compared with either rLCMV-3UTRΔ39-60, rLCMV-UTR-comple, rLCMV-UTR 5-3 change, or rLCMV-Δ26-40 (*P* = 0.0133) (Fig. 4A and B, and Table 4).

**FIG 4.**
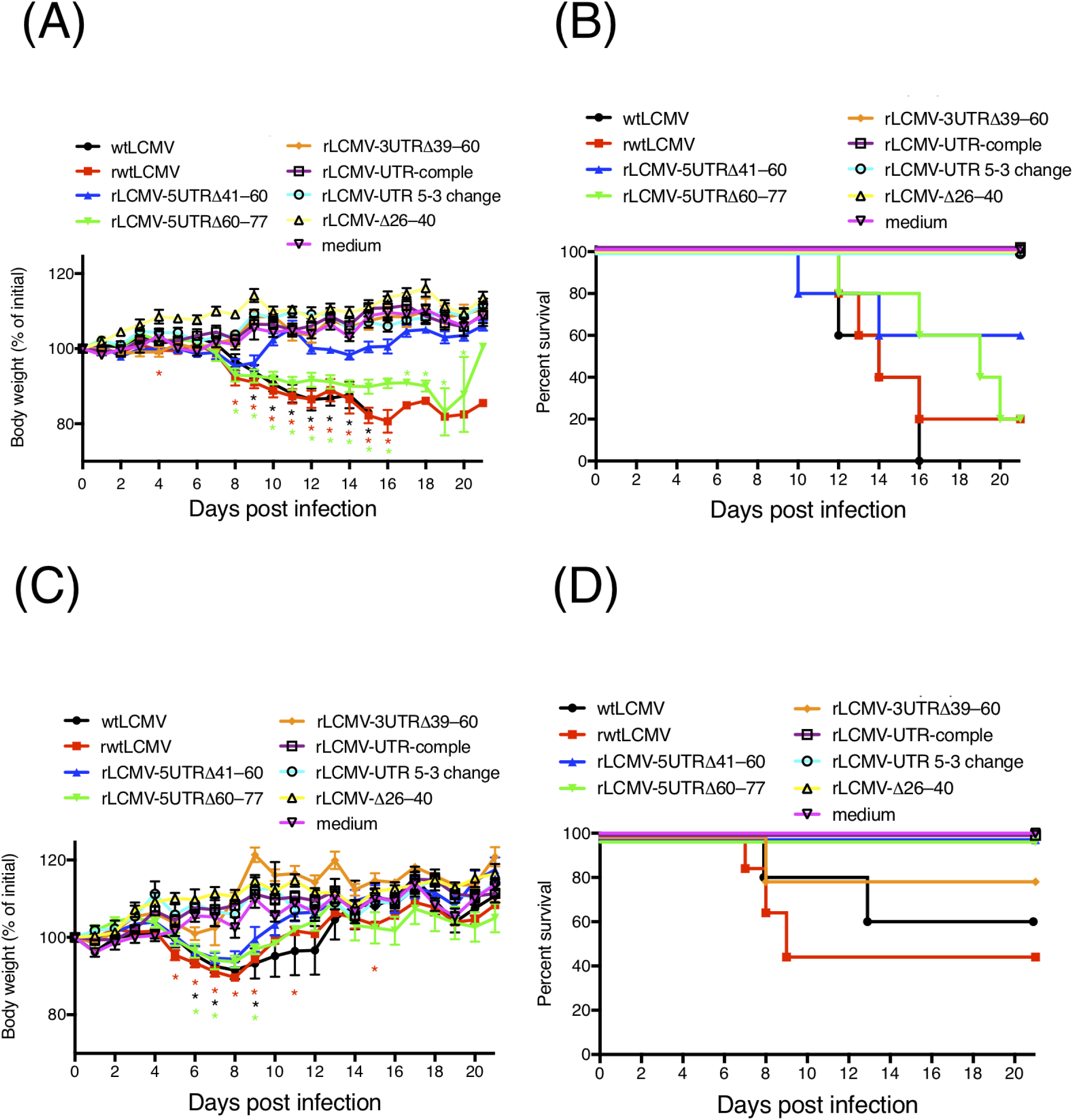
Virulence of wtLCMV and rLCMVs in CBA/NSlc mice and DBA/1JJmsSlc mice. (A and B) Changes in body weight and survival rate of CBA/NSlc mice. Seven-week-old female CBA/NSlc mice (5 mice per group) were intraperitoneally (i.p.) infected with 1.0 × 10^2^ FFU of wtLCMV, rwtLCMV, or various mutated rLCMVs. (C and D) Changes in body weight and survival rate of DBA/1JJmsSlc mice. Eight-week-old female DBA/1JJmsSlc (5 mice per group) were i.p. infected with 1.0 × 10^2^ FFU of wtLCMV or various mutated rLCMVs. Error bars in FIG 4A and C indicate standard errors of the mean. Asterisks indicate that significant differences were observed between the mean body weight of mice infected with medium alone and that of mice infected with LCMVs, displayed in the same colors.

All DBA/1JJmsSlc mice infected with wtLCMV (wtLCMV-DBA/1JJmsSlc mice) or rwtLCMV (rwtLCMV-DBA/1JJmsSlc mice) showed several clinical signs of infection at 5 d.p.i., such as ruffled fur, limb tremors, and weight loss, and approximately half the wtLCMV-DBA/1JJmsSlc mice and rwtLCMV-DBA/1JJmsSlc mice died within 13 d.p.i. (Fig. 4C and D, and Table 4). DBA/1JJmsSlc mice infected with rLCMV-5UTRΔ41-60 (rLCMV-5UTRΔ41-60-DBA/1JJmsSlc mice), rLCMV-5UTRΔ60-77 (rLCMV-5UTRΔ60-77-DBA/1JJmsSlc mice), or rLCMV-3UTRΔ39-60 (rLCMV-3UTRΔ39-60-DBA/1JJmsSlc mice) showed clinical signs of infection by 7 d.p.i. However, limb tremors were not observed and ruffled fur was mild. All rLCMV-5UTRΔ41-60-DBA/1JJmsSlc mice and rLCMV-5UTRΔ60-77-DBA/1JJmsSlc mice showed weight loss but survived. Although one rLCMV-3UTRΔ39-60-DBA/1JJmsSlc mouse showed weight loss and died at 8 d.p.i., 4 out of 5 rLCMV-3UTRΔ39-60-DBA/1JJmsSlc mice did not show weight loss and survived. Significant differences in survival rates were not observed between rwtLCMV and rLCMV-3UTRΔ39-60 (*P* = 0.2194). The hazard ratio for rLCMV / rLCMV-3UTRΔ39-60 was 3.603. All DBA/1JJmsSlc mice infected with either rLCMV-UTR-comple (rLCMV-UTR-comple-DBA/1JJmsSlc mice), rLCMV-UTR 5-3 change (rLCMV-UTR 5-3 change-DBA/1JJmsSlc mice), or rLCMV-Δ26-40 (rLCMV-Δ26-40-DBA/1JJmsSlc mice) showed no clinical signs and survived (Fig. 4C and D, and Table 4).

### Acquired immunity against LCMV in mice induced by infection with rLCMVs

The neutralization titers against LCMV in sera collected from CBA/NSlc mice at 37 days after first infection and DBA/1JJmsSlc mice at 40 days after first infection were evaluated. Among the 53 sera specimens collected from rLCMV-infected mice all specimens but one, collected from an rLCMV-3UTRΔ39-60-CBA/NSlc mouse, showed negative reactions in the focus reduction neutralization test. The 50% focus reduction neutralization titer (FRNT_50_) of the positive sample was 40.

Mice that survived inoculation with rLCMVs were further i.p. infected with 1.0 × 10^3^ FFU of wtLCMV 40 days after the first inoculation. Although all CBA/NSlc mice previously inoculated with medium alone (the control) died within 11 d.p.i., all rLCMV-5UTRΔ41-60-, rLCMV-5UTRΔ60-77-, rLCMV-3UTRΔ39-60-, rLCMV-UTR-comple-, rLCMV-UTR 5-3 change-, and rLCMV-Δ26-40-CBA/NSlc mice survived the secondary viral challenge, exhibiting no clinical signs of infection (Fig. 5A and B, and Table 4).

**FIG 5.**
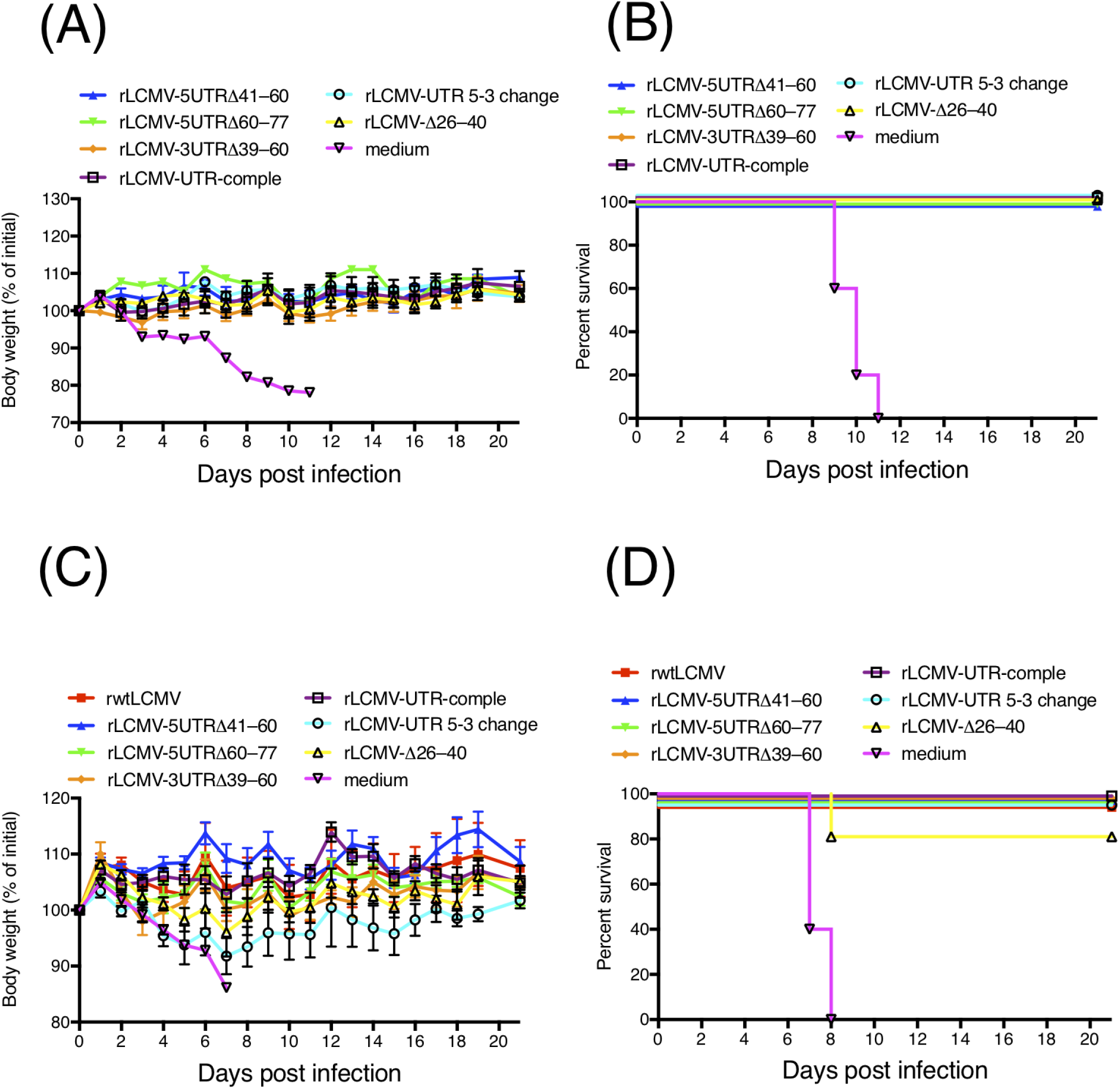
Mice previously infected with rLCMVs were given a lethal dose infection of wtLCMV. Mice that survived inoculation with rLCMVs were further infected intraperitoneally (i.p.) with 1.0 × 10^3^ FFU of wtLCMV 40 days after the first inoculation. (A and B) Changes in body weight and survival rate of CBA/NSlc mice. CBA/NSlc mice previously inoculated with rLCMV-5UTRΔ41-60 (3 mice per group), rLCMV-5UTRΔ60-77 (1 mouse per group), rLCMV-3UTRΔ39-60 (5 mice per group), rLCMV-UTR-comple (5 mice per group), rLCMV-UTR 5-3 change (5 mice per group), rLCMV-Δ26-40 (5 mice per group), or medium (5 mice per group) were used. (C and D) Changes in body weight and survival rate of DBA/1JJmsSlc mice. DBA/1JJmsSlc mice previously inoculated with rwtLCMV (3 mice per group), rLCMV-5UTRΔ41-60 (5 mice per group), rLCMV-5UTRΔ60-77 (5 mouse per group), rLCMV-3UTRΔ39-60 (4 mice per group), rLCMV-UTR-comple (5 mice per group), rLCMV-UTR 5-3 change (5 mice per group), rLCMV-Δ26-40 (5 mice per group), or medium (5 mice per group) were used. Error bars in FIG 5A and C indicate standard errors of the mean.

All control DBA/1JJmsSlc mice died within 8 d.p.i. Conversely, all rLCMV-5UTRΔ41-60-, rLCMV-5UTRΔ60-77-, rLCMV-3UTRΔ39-60-, rLCMV-UTR-comple-, rLCMV-UTR 5-3 change-, and rLCMV-Δ26-40-DBA/1JJmsSlc mice (with the exception of one rLCMV-Δ26-40-DBA/1JJmsSlc mouse) survived the secondary viral challenge (Fig. 5C and D, and Table 4). None of the rLCMV-5UTRΔ41-60-, rLCMV-5UTRΔ60-77-, or rLCMV-3UTRΔ39-60-DBA/1JJmsSlc mice showed any clinical symptoms. One of the 5 rLCMV-UTR-comple-DBA/1JJmsSlc mice showed ruffled fur and weight loss at 6 d.p.i. Several rLCMV-UTR 5-3 change- and rLCMV-Δ26-40-DBA/1JJmsSlc mice showed weight loss 2 d.p.i.; all rLCMV-UTR 5-3 change- and rLCMV-Δ26-40-DBA/1JJmsSlc mice showed ruffled fur 5 d.p.i.; and one rLCMV-Δ26-40-DBA/1JJmsSlc mouse died 8 d.p.i.

### Minigenome assay and VLP assay

To clarify the effect of various mutations or deletions in the UTRs on viral genome transcription and replication, a minigenome assay was established and the function of UTRs with mutated minigenome plasmids (SMGs) was analyzed. To construct SMG-GFP or SMG-luc, cDNA fragments containing the S 5’ UTR, S IGR, GFP or Renilla luciferase open reading frames (ORFs) in an antisense orientation with respect to the 5’ UTR, and the S 3’ UTR were cloned between a murine pol I promoter and a terminator of the pRF vector system (Fig. 6A). The expression of GFP or luciferase was observed only in BHK cells, which were co-transfected with SMG-GFP (or SMG-luc) and both pC-NP and pC-L (Fig. 6B and C). Neither GFP nor luciferase expression was observed in BHK-21 cells transfected with SMG-GFP or -luc and either pC-NP or pC-L.

**FIG 6.**
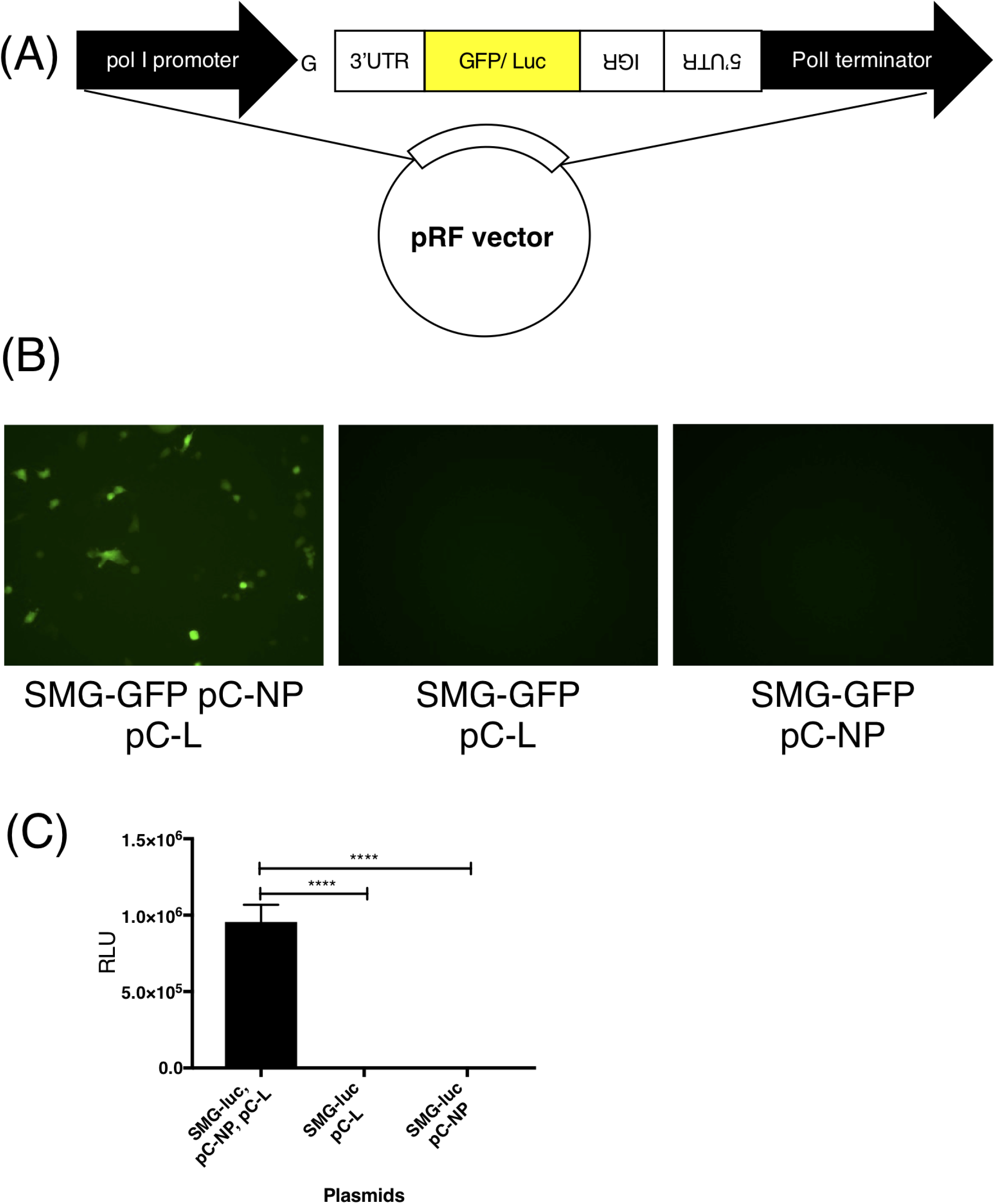
Establishment of a minigenome system for LCMV strain WE (LCMV-WE). (A) Schematic diagram of SMG-GFP or -luc. A cDNA fragment containing the S 5’ UTR, S IGR, GFP or Renilla luciferase ORFs in an antisense orientation with respect to the 5’ UTR, and the S 3’ UTR were cloned between the murine pol I promoter and the terminator of a pRF vector. Additional G residue was inserted between the murine pol I promoter and the viral genome sequence. (B and C) BHK-21 cells were transfected with minigenome plasmids [SMG-GFP (B) or SMG-luc (C)] and both pC-NP and pC-L, or either pC-NP or pC-L. The transfected cells were incubated for 2 days at 37°C then the level of GFP expression was observed under a fluorescent microscope (B) or luciferase activity was measured using the Renilla Luciferase Assay System (C). (\**P* < 0.05, \*\**P* < 0.01, \*\*\**P* < 0.001, \*\*\*\**P* < 0.0001) Error bars indicate standard deviations.

We generated various mutated SMGs in which the mutations were equivalent to the mutations introduced to pRF-WE-SRGs (Table 1 and 2). Detailed information relating to the mutated SMG-GFP or -luc is described in the Materials and Methods section. We evaluated the efficiency of genome transcription and replication of the RNAs derived from SMG-GFP (or -luc), SMG-5UTRΔ20-40-GFP (or -luc), SMG-5UTRΔ41-60-GFP (or -luc), SMG-5UTRΔ60-77-GFP (or -luc), SMG-3UTRΔ20-38-GFP (or -luc), SMG-3UTRΔ39-60-GFP (or -luc), and SMG-UTR-comple-luc (Fig. 7A and B). As previously reported (11), the expression of minigenome-derived reporter genes was hampered by the co-expression of GPC and Z protein (Fig. 7A). Viral genome transcription and replication were either not observed or observed less frequently when cells were transfected with SMG-5UTRΔ20-40-GFP (or -luc). The efficiency of viral genome transcription and replication in cells transfected with SMG-3UTRΔ20-38-GFP (or -luc) or SMG-UTR-comple-luc was significantly lower compared with cells transfected with SMG-GFP (or -luc). The level of luciferase expression in cells transfected with SMG-5UTRΔ41-60-luc, SMG-5UTRΔ60-77-luc, or SMG-3UTRΔ39-60-luc was equivalent to that of cells transfected with SMG-luc (Fig. 7B).

**FIG 7.**
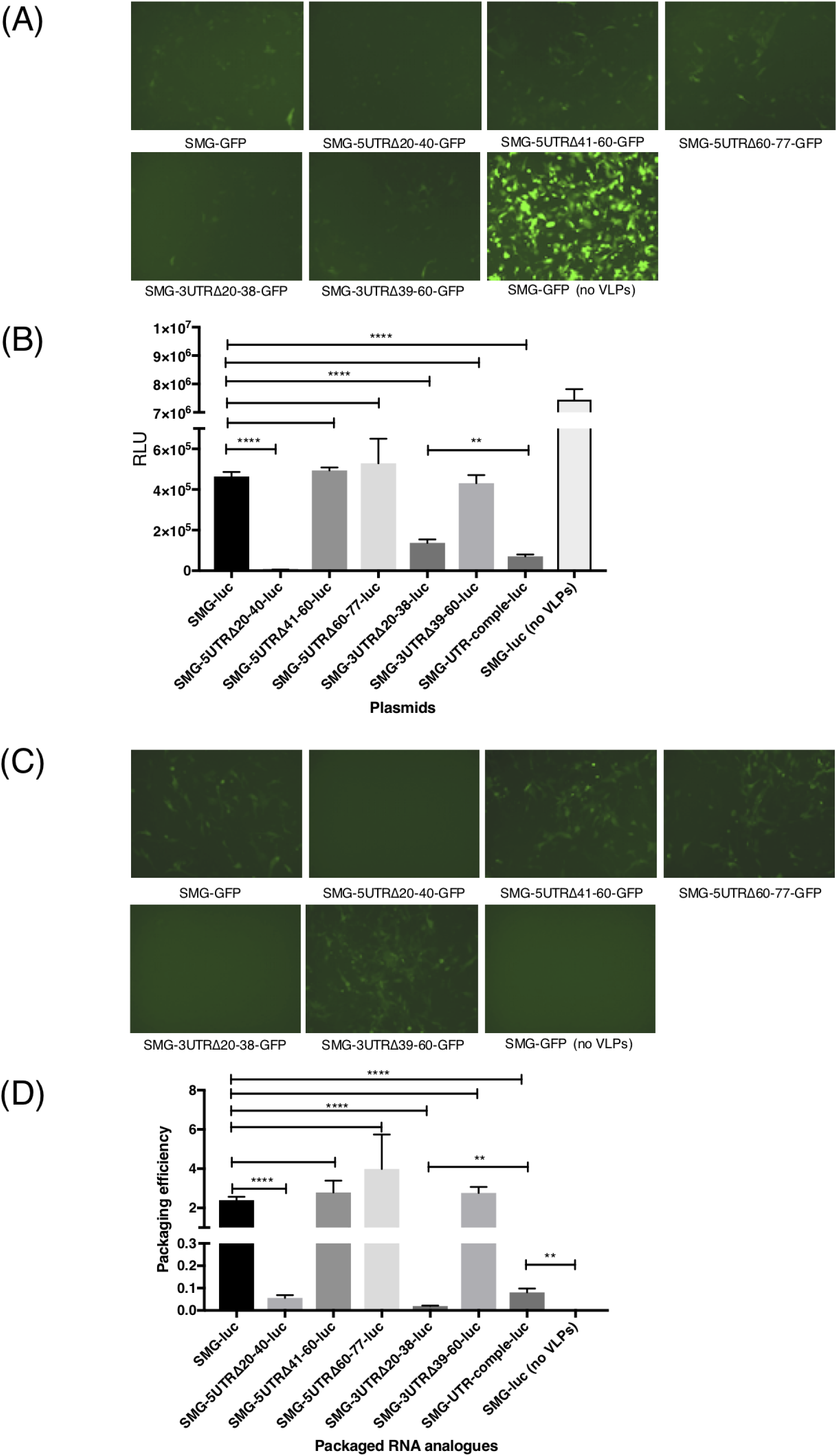
Evaluation of the efficiency of viral genome transcription, replication, and packaging in virus-like particles (VLPs). (A and B) Viral genome transcription and replication were observed in BHK-21 cells transfected with SMG-GFP or -luc, SMG-5UTRΔ41-60-GFP or -luc, SMG-5UTRΔ60-77-GFP or -luc, SMG-3UTRΔ20-38-GFP or -luc, SMG-3UTRΔ39-60-GFP or -luc, or SMG-UTR-comple-luc under conditions of the co-expression of NP, L protein, GPC, and Z protein. BHK-21 cells were transfected with each mutated SMG-GFP or -luc, pC-NP, pC-L, pC-Z, and pC-GPC. As a background control for the rescue of LCMV RNA analogs into VLPs, cells transfected with SMG-GFP or -luc, pC-NP, and pC-L were used [SMG-GFP (no VLPs) and SMG-luc (no VLPs)]. After incubation for 48 hours at 37°C with 5% CO_2_, GFP or luciferase expression was examined. (C and D) Viral genome RNA analogs derived from SMG-GFP or -luc, SMG-5UTRΔ41-60-GFP or -luc, SMG-5UTRΔ60-77-GFP or -luc, SMG-3UTRΔ39-60-GFP or -luc, or SMG-UTR-comple-luc were packaged into VLPs. Supernatants from each well of FIG 7A or B were harvested and used to infect fresh monolayers of BHK-21 cells, which were then incubated for 4 hours at 37°C before adding helper LCMV. Ninety hours post-infection, the passage culture was examined for GFP or luciferase expression. The packaging efficiency of each SMG was determined by dividing the relative light units (RLUs) value obtained from the passage culture 90 hours post-infection by the RLUs value obtained in FIG 7D. (\**P* < 0.05, ** *P* < 0.01, \*\*\**P* < 0.001, \*\*\*\**P* < 0.0001). Error bars indicate standard deviations.

We also evaluated the packaging efficiency of the viral genome RNA analogs into VLPs (Fig. 7C and D). The levels of luciferase expression in cells infected with VLPs which encapsulated RNA products derived from SMG, SMG-5UTRΔ41-60-luc, SMG-5UTRΔ60-77-luc, or SMG-3UTRΔ39-60-luc were approximately equal to each other. Luciferase expression in cells infected with those VLPs that encapsulated RNA products derived from SMG-UTR-comple-luc was significantly lower than that of cells infected with VLPs that encapsulated RNA products derived from SMG-luc, but significantly higher than that of cells treated with supernatant derived from cells transfected with SMG-luc without VLP expression plasmids (pC-Z and pC-GPC) (*P* = 0.0003) (Fig. 7D).

## DISCUSSION

Although there have been limited reports of infection by most arenaviruses, there have been reports of LCMV infection and detection worldwide (1). Furthermore, LCMV has long been widely used not only in virologic research but also in immunological research (9). Understanding the role the LCMV genome plays in virulence and propagation may help to inform the development of new vaccine strategies and/or antiviral therapeutic strategies.

Here we have described how we established an LCMV-WE reverse genetics system and a minigenome system (Fig. 3A and 6). To date, a polymerase-I-driven LCMV-ARM reverse genetics system has been reported by Lukas et al., with analyses of LCMV-ARM using this system also reported (15). The identities of the nucleotide sequences between the S and L segments of LCMV WE and LCMV-ARM are 85% and 82%, respectively. Several reports have described characteristic differences between these strains. Our LCMV-WE reverse genetics system has enabled us to clarify these characteristic differences from the virologic perspective. It has been reported that LCMV-WE infection in non-human primates causes viral hemorrhagic fever that resembles human infection with LASV (26, 27). The LCMV-WE reverse genetics system can enable us to understand the pathogenic mechanisms of viral hemorrhagic fever caused by arenaviruses without the need for Biosafety Level 4 facilities.

The 19 nt of both termini of the genome segments have been reported to be the minimal genomic promoters necessary for viral replication and transcription (29). However, there are dozens of nucleotides in the UTRs of the segments and their function has not yet been studied. In this study, we focused on the S segment UTRs and performed RNA secondary structure prediction of the LCMV S segment UTRs (Fig. 1A). The results of this prediction demonstrated that not only the terminal 19 nt in both 5’- and 3’-termini but also the 28th–33rd nt in the 5’-terminus and the 26th–31st nt in the 3’-terminus formed base pairs with the highest base-pairing probability (Table 3). The results of generating various mutated rLCMVs suggested that the base pairs comprising the 20th–40th nt in the 5’-terminus and the 20th–38th nt in the 3’-terminus, which included the 28th–33rd nt in the 5’-terminus and the 26th–31st nt in the 3’-terminus, greatly affected viral propagation (Fig. 1 to 3, and Table 4). The successful generation of rLCMV-UTR-comple, rLCMV-UTR 5-3 change, and rLCMV-Δ26-40, and the failure to generate rLCMV-Δ20-25, rLCMV-Δ20-30, and rLCMV-Δ31-40 suggested that both the conformation of the panhandle structure and the nucleotide sequences of these base pairs were important for the recognition of the L protein and affected viral propagation. The predicted RNA secondary structure of the rLCMV-Δ26-40 S segment showed no base pairs with high base-pairing probability with the exception of the 19 base pairs in the termini. The reason why rLCMV-Δ26-40 could propagate was not clarified in this study but the conservation of base pairs comprising the 20th–25th nt in both termini and the RNA conformation might enable rLCMV-Δ26-40 to propagate.

The viral growth capacity and focus size of rLCMV-5UTRΔ41-60, rLCMV-5UTRΔ60-77, and rLCMV-3UTRΔ39-60 were equivalent or similar to those of wtLCMV and rwtLCMV (Fig. 3B and D, and Table 4). This also supported the suggestion that the conservation of the base pairs comprising the 20th–40th nt in the 5’-terminus and the 20th–38th nt in the 3’-terminus was not essential but did have a considerable effect on viral propagation, although the other UTRs did not affect viral propagation *in vitro*.

The results of our mice experiments demonstrated that mutations or deletions in UTRs of the S segment of LCMV could attenuate its virulence *in vivo* (Fig. 4, and Table 4). In particular, mutations in the base pairs of the 20th–40th nt in the 5’-terminus and the 20th–38th nt in the 3’-terminus (rLCMV-UTR-comple, rLCMV-UTR 5-3 change, and rLCMV-Δ26-40) caused major attenuation of LCMV *in vivo*. Low efficiency of viral propagation *in vitro* is thought to lead to viral attenuation *in vivo*. Significant differences between the survival curves of rwtLCMV and rLCMV-3UTRΔ39-60 in CBA/NSlc mice and between the survival curves of rwtLCMV and rLCMV-5UTRΔ41-60 or rLCMV-5UTRΔ60-77 in DBA/1JJmsSlc mice were observed. Although there were no significant differences between the survival curves of rwtLCMV and rLCMV-5UTRΔ60-77 or rLCMV-5UTRΔ41-60 in CBA/NSlc mice, or between the survival curves of rwtLCMV and rLCMV-3UTRΔ39-60 in DBA/1JJmsSlc mice, the hazard ratios suggested that rLCMV-5UTRΔ41-60, rLCMV-5UTRΔ60-77, and rLCMV-3UTRΔ39-60 possessed less virulence compared with wtLCMV or rwtLCMV. Recently, several reports have suggested that UTRs in other viruses are involved in virulence. For example, it has been reported that the variable region of the 3’ UTR of the tick-borne encephalitis virus genome was associated with virulence in mice (31, 32). It has also been reported that UTRs of the picornavirus genome can affect viral infection and host innate immunity (33). In this study, the role of the 41st–77th nt in the 5’ UTR and the 39th–60th nt in the 3’ UTR was not elucidated. However, the results of the mice experiments suggested that these UTRs could play a role in affecting host innate immunity *in vivo*. CBA/Nslc mice have been widely used in infectious disease and immunity research (34–36). A single amino acid mutation in Bruton’s tyrosine kinase and reduced function of B cells in CBA/NSlc mice has been reported (37, 38). Although one out of five rLCMV-3UTRΔ39-60-DBA/1JJmsSlc mice lost body weight and died, none of the rLCMV-3UTRΔ39-60-CBA/NSlc mice showed any apparent symptoms and survived the first infection. These results suggest that the 39th–60th nt in the 3’ UTR might affect innate immunity, but further investigation is required to make a firm conclusion about this.

The results of a secondary infection with a lethal dose of wtLCMV in mice that survived suggested that protective immunity was induced in these mice (Fig. 5 and Table 4). rLCMV-UTR-comple-, rLCMV-UTR 5-3 change-, and rLCMV-Δ26-40-DBA/1JJmsSlc mice developed symptoms following the secondary infection and one mouse died at 8 d.p.i. On the other hand, none of the rLCMV-5UTRΔ41-60-, rLCMV-5UTRΔ60-77-, or rLCMV-3UTRΔ39-60-DBA/1JJmsSlc mice showed any clinical symptoms. These results indicate that mice infected with rLCMVs that propagated in a similar way to wtLCMV induce higher protective immunity than mice infected with rLCMVs that had a low capacity for propagation. The results of the neutralization assay suggested that neutralizing antibodies against LCMV were not induced in most mice. This is consistent with previous studies which showed that the generation of anti-LCMV neutralizing antibodies was not detectable for between 60 to 120 days following LCMV-WE infection in mice and that CD8^+^ T-cell-mediated cytotoxicity played a key role in LCMV-WE infection (39).

UTRs play an important role in virus lifecycles, such as viral genome transcription, replication, encapsidation, and packaging into VLPs. It has been reported that deletions in the UTRs attenuated viral growth properties in Bunyamwera virus (BUNV) but that serial passage *in vitro* endowed BUNV with partial recovery of its viral growth properties (40). In that report, the authors found amino acid changes in the C-terminal domain of the L protein and suggested that these changes might be involved in the evolution of the L polymerase, allowing it to recognize the deleted UTRs more efficiently. In our study, the results of a minigenome assay with various mutated MGs suggested that deletions or other mutations in the base pairs comprising the 20th– 40th nt in the 5’-terminus and the 20th–38th nt in the 3’-terminus greatly hampered viral genome transcription and replication (Fig. 7A and B). This seemed to be the cause of the non-production of rLCMVs derived from pRF-WE-SRG-5UTRΔ20-40 and pRF-WE-SRG-3UTRΔ20-38. These results supported the notion that the panhandle structure composed of these base pairs is involved in the recognition site of the L protein. In this study, viruses which acquired mutations in the L protein and recovered their viral growth properties did not emerge in the viral growth-capacity experiment. However, there is a possibility that the L protein of rLCMV-UTR-comple, rLCMV-UTR 5-3 change, and rLCMV-Δ26-40 can undergo amino acid changes and adapt its conformation to the mutated UTRs. Deletions in the other UTRs (the 5’ UTR 41st–60th nt, the 5’ UTR 60th–77th nt, and the 3’ UTR 39th–60th nt) did not affect viral genome transcription and replication efficiency in the minigenome assay or viral genome packaging efficiency in the VLP assay (Fig. 7). These results suggest that UTRs, except for those base pairs composed of 40 nt in the 5’-terminus and 38 nt in the 3’-terminus, were not involved in viral genome transcription, replication, or packaging. Furthermore, these minigenome assay data suggest a low possibility for the conformational adaptation of L protein to the mutated UTRs in rLCMV-5UTRΔ41-60, rLCMV-5UTRΔ60-77, and rLCMV-3UTRΔ39-60.

Although luciferase expression levels from SMG-3UTRΔ20-38-luc were higher than those from SMG-UTR-comple-luc, the packaging efficiency of the RNA products from SMG-UTR-comple-luc was significantly higher compared with that of SMG-3UTRΔ20-38-luc (Fig. 7B and D). These results suggest that RNA products derived from SMG-UTR-comple-luc were certainly packaged into VLPs and carried to the next cells but at low efficiency, and that the sequence and/or conformation of UTRs also affected viral genome packaging efficiency.

In summary, we found that not only the 19 nucleotide base pairs in both termini of the S segment but also the nucleotide base pairs of the 20th–40th nucleotides in the 5’-terminus and the 20th–38th nucleotides in the 3’-terminus of the S segment formed panhandle structures with high base-pairing probabilities. These regions affected viral propagation because they were heavily involved in viral genome transcription and replication in terms of their nucleotide sequence and conformation. Furthermore, our findings suggest that the other UTRs were involved in viral pathogenicity *in vivo*, though they did not affect the efficiency of viral genome transcription, replication, and packaging. The mechanism of attenuation of the virus *in vivo* remains to be determined but the attenuation of LCMV without amino acid changes in component proteins might help us to develop new vaccines.

## MATERIALS AND METHODS

### Cells and viruses

BHK-21 and Vero cells were maintained in Dulbecco’s modified minimal essential medium (DMEM) supplemented with 5% fetal bovine serum (FBS) and 100 μg/ml penicillin– streptomycin (all from Life Technologies, Carlsbad, CA) (DMEM-5FBS) and cultured at 37°C in a 5% CO_2_ atmosphere. LCMV-WE-NIID (GenBank accession numbers LC413283 and LC413284) was amplified in Vero cells and used in this study.

### Immunofocus assay and immunofluorescence assay (IFA)

The infectious dose of LCMV was determined using a viral immunofocus assay. Briefly, after absorption of virus solution into Vero cells cultured in 12-well plates, cells were further cultured for 120 hours at 37°C in DMEM supplemented with 1% FBS and 100 μg/ml penicillin–streptomycin (DMEM-1FBS) with agarose (1%). The cell monolayers were then fixed with 10% formalin in PBS, permeabilized by incubating with 0.2% Triton X-100 in PBS, and stained with anti-LCMV-WE recombinant NP immunized rabbit serum and HRP-goat anti-rabbit IgG (H+L) DS Grd (lot: 917439A, Life Technologies, Carlsbad, CA) (41). Cells were then stained with Peroxidase Stain DAB Kit (Nacalai, Kyoto, Japan) according to the manufacturer’s protocol, and the number of stained foci were counted. For IFA, Alexa Fluor 488 goat anti-rabbit IgG (H + L) (Life Technologies, Carlsbad, CA) was used as the secondary antibody. The cells were observed to determine if they were LCMV-positive or -negative under a fluorescent microscope (BZ-9000, KEYENCE, Osaka, Japan).

### Plasmids

To construct pRF-WE-SRG and pRF-WE-LRG plasmids, cDNA fragments containing either whole S or L segments were cloned between the murine pol I promoter and the terminator of the pRF vector. The pRF vector system was kindly provided by Dr. Shuzo Urata, Nagasaki University and Dr. Juan Carlos de la Torre, of the Scripps Research Institute (San Diego, CA) (17). Insertion of additional G residue directly downstream of the promoter has been reported to enhance the efficiency of both reverse genetics and minigenome systems. The viral cDNA constructs were inserted in sense-orientation for viral complementary (c)RNA. Several mutated pRF-WE-SRG plasmids (pRF-WE-SRG-5UTRΔ20-40, pRF-WE-SRG-5UTRΔ41-60, pRF-WE-SRG-5UTRΔ60-77, pRF-WE-SRG-3UTRΔ20-38, pRF-WE-SRG-3UTRΔ39-60, pRF-WE-SRG-UTR-comple, pRF-WE-SRG-UTR reverse, pRF-WE-SRG-UTR 5-3 change, pRF-WE-SRG-Δ26-40, pRF-WE-SRG-Δ20-25, pRF-WE-SRG-Δ20-30, and pRF-WE-SRG-Δ31-40), which had mutations in the S segment UTRs, were generated using the site-directed mutagenesis method.

To construct SMG-GFP and SMG-luc as minigenome plasmids, cDNA fragments containing the S 5’ UTR, the S IGR, and either GFP or Renilla luciferase ORFs in the antisense orientation to the 5’ UTR, and the S 3’ UTR were cloned between the murine pol I promoter and the terminator of the pRF vector in sense-orientation to the viral cRNA. Additional G residue was also inserted between the murine pol I promoter and the viral minigenome sequence, as well as the reverse genetics system. Several mutated SMG-GFP or -luc plasmids, which had mutations in their S segment UTRs [SMG-5UTRΔ20-40 (-GFP or -luc), SMG-5UTRΔ41-60 (-GFP or -luc), SMG-5UTRΔ60-77 (-GFP or -luc), SMG-3UTRΔ20-38 (-GFP or -luc), SMG-3UTRΔ39-60 (-GFP or –luc), and SMG-UTR-comple (-luc)] were generated using the site-directed mutagenesis method.

The detailed information for each mutated pRF-WE-SRG and SMG-GFP or -luc is shown in Tables 1 and 2. Briefly, pRF-WE-SRG (or SMG)-5UTRΔ20-40, pRF-WE-SRG (or SMG)-5UTRΔ41-60, and pRF-WE-SRG (or SMG) -5UTRΔ60-77 lacked the 20th–40th, the 41st–60th, and the 60th–77th nt residues in the S segment UTRs of their 5’-terminus, respectively. pRF-WE-SRG (or SMG)-3UTRΔ20-38 and pRF-WE-SRG (or SMG)-3UTRΔ39-60 lacked the 20th–38th and the 39th–60th nt residues in the S segment UTRs of their 3’-terminus, respectively. pRF-WE-SRG (or SMG)-UTR-comple had complementary nucleic sequences in the 20th–26th nt residues in their 5’-terminus and the 20th–24th nt residues in their 3’-terminus, respectively. pRF-WE-SRG-UTR 5-3-change had the 20th–38th nt residues in the 3’-terminus in place of the 20th–40th nt residues in the 5’-terminus and vice versa in the 3’-terminus. pRF-WE-SRG-Δ26-40, pRF-WE-SRG-Δ20-25, pRF-WE-SRG-Δ20-30, and pRF-WE-SRG-Δ31-40 lacked the 26th–40th, 20th–25th, 20th–30th, and 31st–40th nt residues in the S segment UTRs of their 5’-terminus, respectively. They also lacked the 24th–38th, 20th–23rd, 20th–28th, and 29th–38th nt residues in the S segment UTRs of their 3’-terminus, respectively.

Plasmids for the expression of LCMV-WE NP, L, Z, and GPC (referred to as pC-NP, pC-L, pC-Z, and pC-GPC, respectively) were obtained as follows. To generate NP, cDNA encoding the NP ORF flanked by EcoRI and NheI restriction sites was amplified by PCR. The PCR product, digested with EcoRI and NheI, was cloned into the EcoRI-NheI restriction site of plasmid pCAGGS. To generate pC-L, the L ORF flanked by KpnI and NheI restriction sites was amplified by PCR. The PCR product was cloned into the KpnI-NheI restriction site of pCAGGS. To generate Z, cDNA encoding the Z ORF flanked by EcoRI and NheI restriction sites was amplified by PCR. The PCR product, digested with EcoRI and NheI, was cloned into the EcoRI-NheI restriction site of plasmid pCAGGS. To generate pC-GPC, the GPC ORF flanked by EcoRI and NheI restriction sites was amplified by PCR. The PCR product was cloned into the EcoRI-NheI restriction site of pCAGGS.

### Transfection, minigenome, and reverse genetics system

For the minigenome, 1.0 × 10^5^ BHK-21 cells were seeded into 24-well plates to reach approximately 80% confluence. Cells were transfected with minigenome plasmids (SMG-GFP or SMG-luc), pC-NP, and pC-L using TansIT-LT1 DNA transfection reagent (Mirus Bio, Madison, WI). The total amount of DNA transfected ranged from 0.8 μg to 2.6 μg depending on the volume of pC-NP and pC-L added, but the amount of transfection reagent was kept constant at three volumes (μl) per added DNA (μg). The transfected cells were incubated for 2 days at 37°C, then the GFP expression level was observed under a fluorescent microscope and luciferase activity was measured using a Renilla Luciferase Assay System (Promega, Fitchburg, WI). The detailed methodology is described in the luciferase assay section.

For the reverse genetics system, 3.0 × 10^5^ BHK-21 cells were seeded into 6-well plates to reach approximately 80% confluence. Cells were transfected using TansIT-LT1 DNA transfection reagent with 0.8 μg pRF-WE-SRG, 1.4 μg pRF-WE-LRG, 0.8 μg pC-NP, and 1.0 μg pC-L. The total amount of DNA transfected was 4.0 μg and the amount of transfection reagent used was 12.0 μl. Under these transfection conditions, the cells were incubated for at least 3 days at 37°C. The supernatant was harvested and the infectious dose of recombinant virus was measured using an immunofocus assay or IFA. The RNA of each rLCMV was extracted and viral genome sequences of all rLCMVs were confirmed to be as intended using the Sanger Sequence method.

### Rescue of LCMV RNA analogs into LCMV-like particles

The rescue of LCMV RNA analogs into VLPs was carried out as previously reported (11), with 3.0 × 10^5^ BHK-21 cells in 2.0 ml of DMEM-5FBS seeded into 6-well plates to reach approximately 80% confluence. The cell culture medium was changed to DMEM supplemented with 2% FBS and 100 μg/ml penicillin–streptomycin (DMEM-2FBS). The cells were transfected using TansIT-LT1 DNA transfection reagent with 1.0 μg SMGs-GFP or -luc (SMG-GFP or -luc or various mutated SMGs-GFP or -luc), 0.8 μg pC-NP, 1.0 μg pC-L, 0.1 μg pC-Z, and 0.3 μg pC-GPC. As a background control for the rescue of LCMV RNA analogs into VLPs, cells transfected with 1.0 μg SMG, 0.8 μg pC-NP, and 1.0 μg pC-L were used [SMG-GFP (no VLPs) and SMG-luc (no VLPs)]. The amount of transfection reagent was kept to three volumes (μl) per amount of DNA added (μg). After incubation for 48 hours at 37°C with 5% CO_2_, the expression of GFP or luciferase was examined, then 1.2 ml of supernatant from each well was harvested. Fresh monolayers of BHK-21 cells seeded into 6-well plates were infected with the supernatants. Cells were incubated with the supernatants for 4 hours at 37°C and infected with helper LCMV at a multiplicity of infection (moi) of 2 FFU/cell. Ninety hours post-infection, the cells were examined for GFP or luciferase expression.

### RNA secondary structure prediction

To predict RNA secondary structures the web application CENTROIDFOLD was used (http://www.ncrna.org/centroidfold/) (42). The RNA sequences of the 5’ and 3’-termini UTRs and 50 nt lengths of ORF (GPC or NP ORF) regional RNA sequences, which were directly downstream of the UTRs, were linked and sent to the CENTROIDFOLD server. The CONTRAfold model (weight of base pairs: 2^2) was used to calculate base-pairing probabilities. The results of RNA secondary structure prediction were referenced in the generation of rLCMVs, in which mutations were introduced into the UTRs.

### Luciferase assay

Cells were washed with PBS and lysed with 100 μl Renilla Luciferase Assay System Lysis Buffer (Promega). Cell lysates (20μl) were mixed with 100 μl Renilla Luciferase Assay System Substrate (Promega). The luminescence level was measured and the relative light units (RLUs) of luciferase were determined using GloMax 96 luminometer (Promega) according to the manufacturer’s protocol. To analyze the efficiency of viral genome packaging, the RLUs acquired from cells infected with VLPs was divided by the RLUs acquired from cells transfected with corresponding minigenome plasmids. Bar graphs were drawn using GraphPad Prism software (GraphPad Software, Inc.) and statistically analyzed using an unpaired two-tailed *t*-test.

### Viral growth kinetics

Viral growth kinetics of wtLCMV, rwtLCMV, and recombinant LCMV with various mutations in Vero cells were analyzed. Briefly, confluent monolayers of Vero cells cultured in 12-well plates were infected with each LCMV at an moi of 0.01 per cell. Cells were washed 3 times with DMEM-2FBS after a one-hour adsorption period at 37°C, and 1 ml DMEM-2FBS was added to each well. Supernatant samples were collected at 0, 24, 48, 72, 96, and 120 h post-infection. The supernatants were centrifuged at 8,000 g for 5 min to remove cell debris and stored at −80°C until the infectious dose was measured using a viral immunofocus assay. Viral growth curves of LCMVs were drawn using GraphPad Prism software (GraphPad Software, Inc.) and statistically analyzed using two-way ANOVA.

### Animal experiments

Animal experiments were performed under Animal Biosafety Level 3 laboratory. All experiments were performed in accordance with the Guidelines for Animal Experimentation of National Institute of Infectious Diseases (NIID), Japan, under the approval of the Committee on Experimental Animals at NIID (No. 116038). Specific-pathogen-free 7-week-old female CBA/NSlc mice and specific-pathogen-free 8-week-old female DBA/1JJmsSlc mice were purchased from Japan SLC, Inc. (Shizuoka, Japan). Mice were i.p. infected with 1.0 × 10^2^ FFU wtLCMV, rwtLCMV, mutated rLCMVs, or medium alone. Mice were monitored daily for 21 days for clinical symptoms, body weight, and survival. Mice showing more than 20% weight loss were euthanized out of ethical consideration. Blood was collected from the caudal vein of each mouse that survived to 37 days post-infection (CBA/NSlc mice) or 40 days post-infection (DBA/1JJmsSlc mice). Following the first infection, mice that survived and showed no apparent symptoms at 40 days post-infection were further i.p. inoculated with 1.0 × 10^3^ FFU of wtLCMV and again monitored daily for 21 days for clinical symptoms, body weight, and survival. After 21 days post-infection, mice were euthanized under isoflurane deep anesthesia. Survival curves were drawn using GraphPad Prism software (GraphPad Software, Inc.) and statistically analyzed using the log-rank (Mantel–Cox) test. The curves of body weight changes were drawn using GraphPad Prism software (GraphPad Software, Inc.) and statistically analyzed using multiple t-tests. Discovery was determined using the two-stage linear step-up produced by Benjamini, Krieger, and Yekutieli, with Q = 5% (43).

### Neutralization assay

Blood was collected from the caudal vein of each mouse at either 37 d.p.i. or 40 d.p.i. using BD Microtainer blood collection tubes (BD, Franklin Lakes, NJ) and centrifuged at 8,000 g for 5 min. Separated plasma was inactivated by heat-treatment at 56°C for 30 min. Plasma was serially diluted with DMEM-2FBS from 1:20 to 1:160 (two-fold serial dilution) and mixed with an equal volume of DMEM-2FBS containing 40–70 FFU/50 μl wtLCMV. The mixtures were incubated for 1 hour at 37°C. Vero cells cultured in 12-well plates were inoculated with 100 μl of each mixture and cultured at 37°C for 1 hour for adsorption. The cells were overlaid with 1 ml maintenance medium (Eagle’s minimal essential medium containing 1% methylcellulose, 2mM L-glutamine, 0.22% sodium bicarbonate, and 2% FBS). The plates were incubated at 37°C in 5% CO_2_ for 120 hours. Cells were fixed with 10% formaldehyde, permeabilized by incubation with PBS containing 0.2% Triton X-100 (SIGMA-ALDRICH, St. Louis, MO), and then stained with anti-LCMV-WE recombinant NP immunized rabbit serum and HRP-goat anti-rabbit IgG (H+L) DS Grd (Life Technologies, Carlsbad, CA) (41). The cells were then stained with Peroxidase Stain DAB Kit (Nacalai, Kyoto, Japan) according to the manufacturer’s protocol. The number of stained foci were counted as described above, and the FRNT_50_ was measured. The FRNT_50_ titers were defined as the reciprocal of the serum dilution level at which the focus number became less than 50% of the control, the focus number of the wells inoculated with the mixture of non-plasma containing DMEM-2FBS and DMEM-2FBS containing 40–70 FFU/50 μl of wtLCMV.

## ACKNOWLEDGMENTS

Japan Society for the Promotion of Science (JSPS) provided funding to Satoshi Taniguchi under grant numbers 15K21645 and 11J02534. JSPS provided funding to Masayuki Shimojima under grant number 16K08041. Japan Agency for Medical Research and Development (AMED) provided funding to Masayuki Saijo under grant numbers 19fk0108072j0002.

We thank Prof. Juan Carlos de la Torre and Dr. Shuzo Urata for providing us with the pRF vector system and giving us technical advice on how to rescue recombinant LCMV. We thank Prof. Yoshiharu Matsuura for providing us with LCMV-WE-NIID strain. We thank Dr. Koichiro Iha, Dr. Kie Yamamoto, Dr. Yoshimi Tsuda, Ms. Momoko Ogata, Dr. Yuto Suda, Ms. Makiko Ikeda, Mr. Kenichi Shibasaki, and Ms. Nor Azila Muhammad Azami for their technical assistance. This research was supported by Grant-in-aids from the Japan Society for the Promotion of Science KAKENHI (15K21645, 11J02534, and 16K08041).

